# Genome-wide characterization of genetic and functional dysregulation in autism spectrum disorder

**DOI:** 10.1101/057828

**Authors:** Arjun Krishnan, Ran Zhang, Victoria Yao, Chandra L. Theesfeld, Aaron K. Wong, Alicja Tadych, Natalia Volfovsky, Alan Packer, Alex Lash, Olga G. Troyanskaya

**Affiliations:** Lewis-Sigler Institute for Integrative Genomics, Princeton University, Princeton, NJ 08544; Department of Molecular Biology, Princeton University, Princeton, NJ 08544; Department of Computer Science, Princeton University, Princeton, NJ 08544; Simons Center for Data Analysis,Simons Foundation, NY 10010 New York

## Abstract

Autism spectrum disorder (ASD) is a range of major neurodevelopmental disabilities with a strong genetic basis. Yet, owing to extensive genetic heterogeneity, multiple modes of inheritance and limited study sizes, sequencing and quantitative genetics approaches have had limited success in characterizing the complex genetics of ASD. Currently, only a small fraction of potentially causal genes—about 65 genes out of an estimated several hundred—are known based on strong genetic evidence. Hence, there isa critical need for complementary approaches to further characterize the genetic basis of ASD, enabling development of better screening and therapeutics. Here, we use a machine-learning approach based on a human brain-specific functional gene interaction network to present a genome-wide prediction of autism-associated genes, including hundreds of candidate genes for which there is minimal or no prior genetic evidence. Our approach is validated in an independent case-control sequencing study of approximately 2,500families. Leveraging these genome-wide predictions and the brain-specificnetwork, we demonstrate that the large set of ASD genes converges on a smaller number of key cellular pathways and specific developmental stages of the brain. Specifically, integration with spatiotemporal transcriptome expression data implicates early fetal and midfetal stages of the developing human brain in ASD etiology. Likewise, analysis of the connectivity of top autism genes in the brain-specific interaction network reveals the breadthof autism-associated functional modules, processes, and pathways in the brain. Finally, we identify likely pathogenic genes within the most frequent autism-associated copy-number-variants (CNVs) and propose genes and pathways that are likely mediators of autism across multiple CNVs. All the predictions, interactions, and functional insights from this work are available to biomedical researchers at asd.princeton.edu.

## Introduction

Complex human diseases such as autism are driven by a multitude of genetic variants across the genome that manifest as a range of developmental and functional perturbations, often in specific tissues and cell types^1, 2^. Sequencing-based discovery efforts^3–    8^ have produced valuable catalogues of genetic variants that point toward potential causal genes of autism. In particular, recent large exome sequencing studies of autism simplex families, as well as case-control cohorts, have combined the evidence from both *de novo* and transmitted loss-of-function mutations to implicate around 65 genes in autism risk^9^. However, this represents only a fraction ofthe estimated 400-1,000 genes likely involved in autism susceptibility^10, 11^. To discover the complete genetic basis of autism and identify the dysregulated cellular and developmental functions of the brain underlying the disease, we need approaches that leverage previously implicated genes in the context of brain-specific biology to discover the full complement of autism-associated genes.

Molecular interaction networks are effective summaries of our knowledge about cellular processes and powerful computational tools for investigating disease genes complementary to sequencing or genetic studies. Previous network-based analyses that focused on autism^12–         22^ (reviewed in^23^)demonstrate the promise of functional genomics approaches. These methods, however, are limited by the underlying networks themselves. Commonly used networks are based on sets of genes or proteins that are co-expressed in asingle developmental context or on protein physical interaction networks that lack tissue-specificity and are still largely incomplete in human. Existing methods, while invaluable, are nevertheless focused on known risk genes or genes with strong prior genetic evidence of ASD association, thus limiting the potential for discovering the full spectrum of disease associated genes.

We address these challenges with an evidence-weighted machine learning approach that utilizes a brain-specific functional interaction network^24^. The brain-specific network was built by integrating thousands of genomic experiments to create a genome-wide probabilistic graph representing how genes function together in pathways in the brain, or, intuitively, a molecular-level functional map of the brain^24^. The evidence-weighted disease-gene classifier, described here, learns the connectivity patterns of known ASD genes in this brain network and then uses these data-driven signals specific to ASD-associated genes to predict the level of potential ASD association for every gene in the genome. Using this approach, here we provide predictions of ASD-associated genes, including candidate genes that have minimal or no prior genetic evidence. Many of these candidate genes have been validated by sequencing studies as *bona fide* ASD-associated genes since the initial predictions were made.Through integrated analysis of the network and our top predictions, we identify developmental stages implicated in ASD, characterize frequent autism-associated CNVs, and discover functional modules potentially dysregulated in the autistic brain. The genome-wide complement of autism candidate genes produced in this study can then be used to systematically prioritize genes for re-sequencing and validation through genetic association, as well as guide the analysis of whole genome sequencing results. This has the potential to accelerate discovery of the full genetic spectrum underlying ASD needed to refine genetic diagnosis and develop treatments.

## Results

We use a human brain-specific functional interaction network (developed recently^24^) to predict ASD candidate genes across the genome and then use these predictions to systematically explore the developmental and functional features of the molecular phenotype of autism (Figure 1). Our predictions provide biomedical researchers with a diverse set of ASD candidate genes that they can explore in the context of the underlying brain functional network to further understand autism genetics and develop more focused genetic screening toolkits.

**Figure 1:**
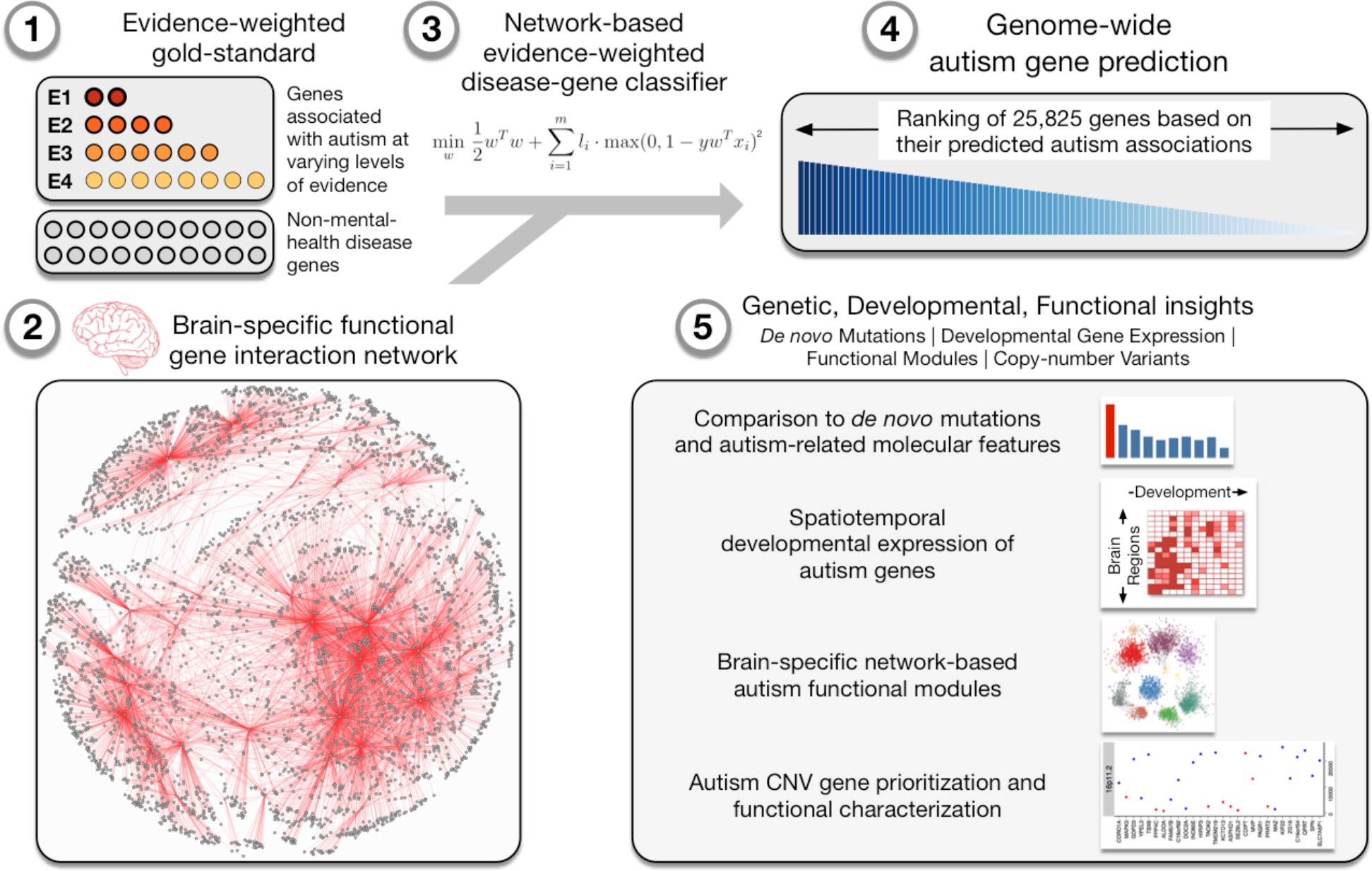
Genome-wide prediction of autism-associated genes. Our ASD-gene predictions are based on a machine learning approach that (1) uses a gold-standard of known disease genes-those linked to autism with varying levels of evidence (E1-4) as positives, and other genes linked to non-mental-health diseases as negatives-in the context of a (2) human brain-specific functional interaction network to (3) build an evidence-weighted network-based classifier capturing autism-specific gene interaction patterns, and (4) predict the probability of autism association of each gene across the genome. We demonstrate the accuracy and utility of our genome-wide complement of autism-associated genes by (5) validating these predictions with *de novo* autism-associated mutations from an independent sequencing study; elucidating the spatiotemporal developmental gene-expression patterns of top-ranked autism genes; laying out the landscape of autism-associated brain-specific functional modules (network clusters); and prioritizing candidate causal genes within large intervals of recurrent autism-associated copy-number variants.

### Genome-wide prediction of autism-associated genes

We recently developed a gene-interaction network model containing predicted functional relationships for all pairs within 25,825 genes in the human genome specifically in the context of human tissues, including the brain^24^. The brain-specific network was built using a Bayesian method that extracts and integrates brain-specific functional signals from thousands of gene expression, protein-protein interaction, and regulatory-sequence datasets. Intuitively, the Bayesian approach first learns a “weight” for each input dataset based on how relevant and accurate it is in reflecting how cellular pathways function in the brain. Then the method predicts a probability of functional relationship, reflecting how likely each pair of genes in the genome is to participate in the same pathway in the brain, calculated by probabilistically weighted integration of relevant signals across input datasets (see Supplemental Note 1 for more details).

We developed an evidence-weighted network-based machine-learning method that uses this brain-specific network to systematically discover novel candidate ASD-associated genes across the genome. We first curated 594 genes linked with autism from a number of publicly available databases prior to 2014, ranging from high-confidence genetic associations (e.g., SFARI Gene^25^ and OMIM^26^) to automatically text-mined frequent ASD-gene co-occurrences in published abstracts (e.g., Gene2Mesh (gene2mesh.ncibi.org) and DGA^27^). We grouped these genes into four evidence levels based on the strength of evidence associating them with ASD (E1-4; Table S1). Using these genes along with their evidence levels as positive gold-standard examples and genes annotated to non-mental-health diseases in OMIM as negative examples, we trained an evidence-weighted Support Vector Machine (SVM) classifier using the connectivity of these gold-standard genes to all the genes in the human brain-specific network as features (Figure 1; see *Methods*). Intuitively, this approach first identifies network patterns that characterize known ASD-related genes while taking into account the level of “trust” in each gene’s association with autism. The patterns consist of genes in the network that are differentially (strongly or weakly) connected to known ASD genes (positives) relative to genes linked to non-mental-health diseases (negatives). The classifier then uses these patterns to calculate a score of potential ASD association for each gene in the genome, identifying novel ASD candidate genes as those that highly resemble known ASD genes in their interaction pattern in the network (Figure S1; Table S2). This produces a comprehensive, genome-wide ranked list of autism candidate genes (Table S3) that is robust to variations in our training gold standard (Figure S2). To improve interpretability of these ASD gene predictions, we estimated probabilities (using isotonic regression) and permutation-based *P* values (and corresponding FDR *Q* values) for each gene (Figure S3, see *Methods*).

Evaluating only on held-out high-confidence (E1) genes through five-fold cross-validation (repeatedly training a model on 4/5^th^ of E1 genes and testing on the hidden 1/5^th^), we found that our approach was accurate (AUC = 0.80; Figure 2a). Moreover, this evidence-weighted classifier trained using E1-4significantly outperformed an unweighted classifier trained using only high-confidence E1 genes (AUC = 0.73), a weighted classifier using only E1 and E2 (AUC = 0.76), and a weighted classifier using E1 and E2 supplemented with random genes that match E3 and E4 in annotation, expression, and gene-length characteristics (AUC = 0.74), thus unequivocally establishing that including lower-confidence genes (E3 and E4) in an evidence-weighted framework helps in significantly improving performance (Figure 2a). Intuitively, this demonstrates that although the individual genes in these lower confidence sets (E3 and E4) are not validated (a substantial fraction may not end up being associated with ASD pathogenesis), as a unit, they still contain signal that is informative for ASD. Our computational approach effectively leverages these signals to improve ASD gene prediction performance.

**Figure 2:**
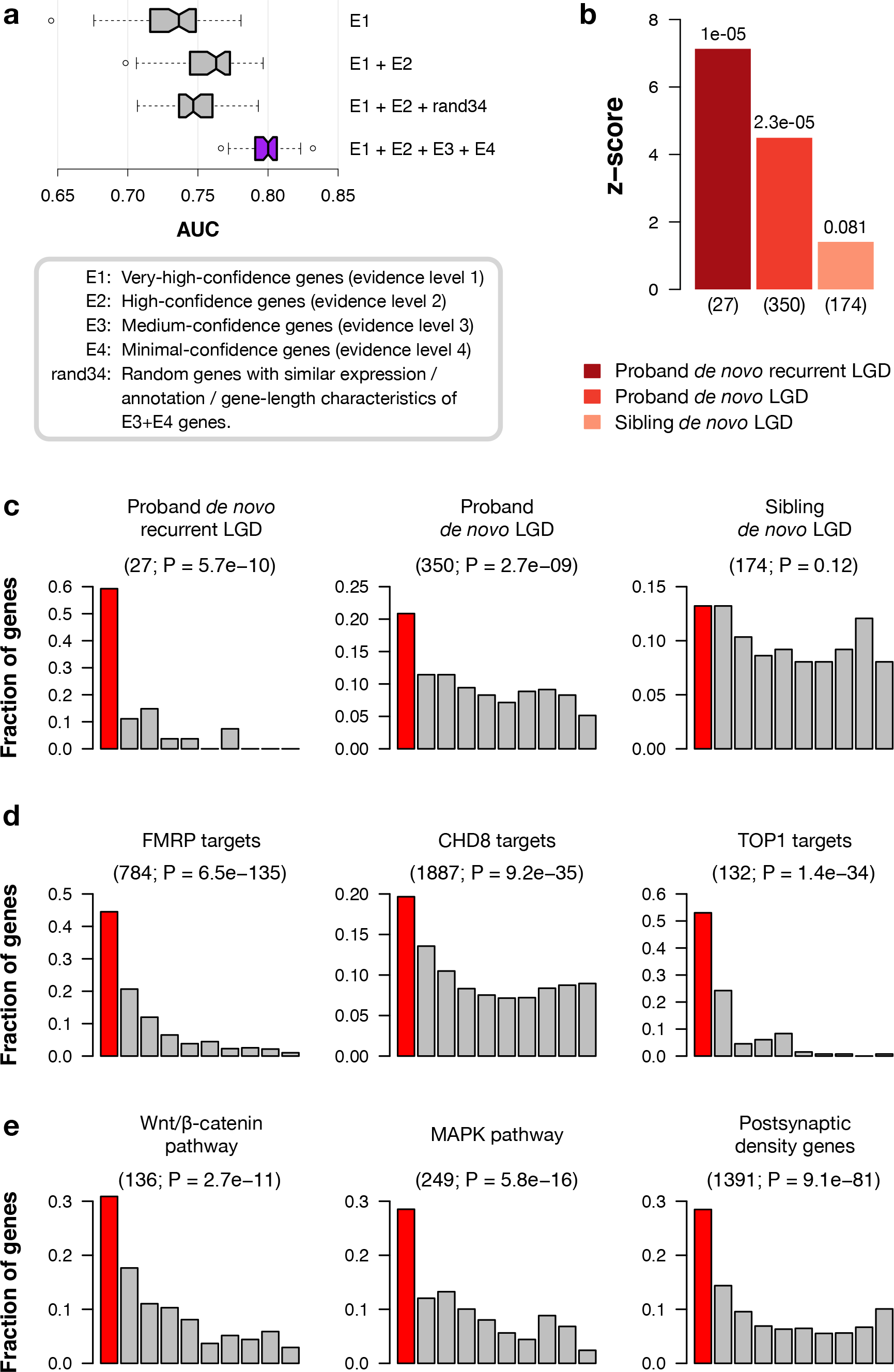
Evaluation of autism-gene predictions. (**a**) Plot of area under the ROC curve (AUC) of autism gene prediction with different versions of training gold-standards. Including lower confidence genes in an evidence-weighted framework helps in predicting autism genes, whereas replacing them with a random set of genes with matched expression, annotation, and gene length characteristics (rand34) decreases performance. The evidence-weighted classifier (purple) significantly outperforms all the other classifiers (Wilcoxon rank-sum test; *P*<1e-14). (**b**) Rank-based enrichment test on mutation data from an independent sequencing study (SSC 2014). The plot shows the z-score quantifying the enrichment of the three gene-sets of interest (labeled in the legend below) towards the top of our genome-wide ranking of genes (see *Methods*). The number of genes in the gene-sets is given in parenthesis below each bar. Our top-ranked genes are significantly enriched in gene targets of *de novo* likely-gene-disrupting (LGD; also known as loss-of-function) mutations observed in probands (autistic children; second bar). This enrichment is striking for targets of recurrent LGD mutations in probands (first bar), and diminishes for LGD-targets in unaffected siblings of the autistic probands (third bar). The *P* values recorded at top of each bar were calculated using a permutation test described in *Methods* in detail. (**c-d**) All evaluations presented here were carried out by dividing our genome-wide predictions into deciles and using a one-sided binomial test to calculate the significance of a specific gene-set of interest among the first decile (top 10%) of the predictions. Each bar plot shows the fraction of genes in the gene-set (y-axis) that occur within each of the 10 deciles (x-axis), with the first decile colored in red. The name of the gene-set is given on top of each plot along with the total number of genes in the gene-set and the *P* value from the binomial test in parenthesis just below. (c) Mutation data from the 2014 SSC study (used in b) evaluated within the top 10% of our predictions shows a trend-significant enrichment of proband LGD mutations (first and second plots) and non-enrichment of sibling LGD mutations (third plot)-that is consistent with those observed with rank-based test (shown in b). (**d**) Targets of major ASD-associated regulatory proteins: Experimentally determined gene targets of FMRP, CHD8 and TOP1 are also significantly enriched in the top 10% of our predictions. (**e**) Members of major ASD-associated pathways/complexes: Similar significant enrichment was also observed for genes involved in Wnt signaling, MAPK signaling, and the postsynaptic density complex.

In addition to computational evaluation by cross-validation, we performed a systematic empirical evaluation based on results from an external exome-sequencing study of 2,517 families^7^. We focused on *de novo* likely-gene-disrupting (LGD; also known as loss-of-function) mutations identified in these families with one autistic child (proband), and, in most cases, an unaffected sibling (Table S4). Genes harboring LGD mutations in probands are significantly enriched towards the top of our ranking independent of any specific cutoff (permutation test *P* = 2.3e-5, see *Methods);* this enrichment is even more pronounced for recurrent proband LGDs (mutations found more than once among autistic children), which are highly likely to be true autism-associated genes (P = 1e-5). In contrast, this enrichment is absent for genes with LGD mutations in unaffected siblings (P = 0.081), showing that our predictions are specifically ranking LGD-target genes related to autism over those potentially unrelated to the disease (Figure 2b). For further analysis, we focus on the top decile of predicted ASD-associated genes (median FDR<0.025), which reflects the above trend: these ∼2,500 genes are significantly enriched for LGD mutations in probands (binomial test *P* = 2.7e-9; Figure 2c; see *Methods*), but not in unaffected siblings (*P* = 0.12). The enrichment is even more striking for proband recurrent LGDs (*P* = 5.7e-10),with about 60% of these proband recurrent LGDs present in the top decile. It is important to note that our prediction method does not use the quantitative genetics data as input, and predictions for each gene are made through cross-validation where any potential information about that gene’s known connection to autism (its presence in E1-E4) is “hidden” from the method. In fact, the trends observed in this analysis hold even when the evaluation standard is strictly limited only to ‘novel’ genes not used in our training set that were observed in families still unpublished at the time of our analysis (Figure S4) or when tested on genes identified in a study independent of the SSC^8^ (*P* = 4.12e-6). These trends are also not non-specifically biased by long, brain-expressed genes (Supplemental Notes 2 and 3; see *Methods*). In contrast to prior network-based methods, our approach is able to identify novel candidate genes with no prior genetic link to ASD (Figure S5), while maintaining accuracy (Figures S6).

Our top-ranked first decile predictions include a significant number of experimentally identified targets of major ASD-associated regulators (FMRP^28^, TOP1^29^, and CHD8^30^), as well as members of some of the major pathways implicated in autism (Wnt/b-catenin^4, 21^, MAPK^31^, post-synaptic density^32^) (Figure 2c,d). Furthermore, we found that interaction partners of brain-expressed isoforms of known ASD genes identified in an independent, high-throughput protein interaction assay^33^ were also enriched in the first decile of our ranking (binomial test *P* = 1.96e-5). Our top-ranked genes are specific to autism, exhibiting an expected overlap with genetically similar disorders (e.g., intellectual disability and schizophrenia), while being distinct from genes associated with other unrelated neuronal diseases (e.g., Alzheimer’s and Parkinson’s diseases; Figure S7). Further, in agreement with recent findings^34^, genes we predict to be highly linked to ASD tend to be more evolutionarily constrained than all the other genes (permutation test *P* = 1e-6; Figure S8, Supplemental Note 4; see *Methods*).

Our predictions (Table S3) contain genes with prior ASD-association that were not part of our positive training set. Variants in GNAO1 (rank #3)-encoding G-proteins that couple to receptors which function in the central nervous system-give rise to motor defects with associated behavioral phenotypes^35^ and can cause involuntary movements as well as severe developmental delay^36^. The gene SYN2 (rank #7) has replicated evidence for maternally-inherited frame-shift and missense mutations associated with male ASD cases^37^, and Syn2 deleted mice show impaired social behavior and memory, as well as abnormal exploration of novel environments^38^. ATXN1 (rank #8) is deleted in five out of six clinical cases, most of whom had ASD, developmental speech delay, and ADHD^39^. Two independent cases in the Simons Simplex Collection have been identified to harbor *de novo* nonsense mutations in the neuronal ankyrin gene ANK2^6, 40^ (rank #9). We highlight many more examples of such top-ranked genes in Supplemental Note 5.

With the results above validating our ranking, our chief contribution is the prediction of several genes with completely novel associations to autism. Interestingly, many among these novel top-decile genes are within large CNV intervals reported in autism cases, potentially implying their role in the pathogenicity of those CNVs. SYT1 (rank #5), for example, is a synaptic membrane protein in the CNV 12q21.2-21.31, which has been observed to be duplicated or deleted in autism cases in two large case-control studies^41, 42^. A largely uncharacterized gene, OLFM1 (rank #14) is within 9q33.2-34.4, a CNV with 5 duplications reported in autism cases^42^. Similarly, NOVA1 (rank #18) resides in the CNV 14q 11.2-21.1; two cases with duplication of this CNV are found to suffer from developmental delay, intellectual disability, or ASD^42^. As potential drivers within CNVs, such genes are prime candidates for follow-up with experimental verification and genetic validation.

Taken together, these rigorous evaluations based on several large, independent, experimentally derived data sets, along with evidence from published literature for individual genes, demonstrate that our genome-wide predictions capture a major molecular signature of autism. Our predictions can therefore be used for systematic analyses to further insights into autism genetics and pathogenesis.

### Autism-linked genetic changes in fetal brain development

One of the crucial characteristics of autism is altered brain development^43^. Identifying the precise developmental stages and regions that are affected by the genetic changes underlying autism is crucial in unraveling the disease mechanism. Here, we use our genome-wide ASD-associated gene predictions in concert with large-scale spatiotemporal gene-expression data of the developing human brain^44^ to identify regions and stages most pertinent to autism. Previous computational studies have used these data to construct coexpression networks^17, 40^ bypooling multiple regions or developmental stages to obtain a minimum number of samples for coexpression analysis. Our approach leverages the comprehensive set of ASD candidate genes and thus, for the first time, allows analysis of individual regions and time-points within these data in the context of autism. We tested the enrichment of gene expression signatures specific to each spatiotemporal window in our autism-gene predictions using a carefully controlled permutation test to identify when and where predicted ASD genes are specifically active (Figure 3, Table S5, Figure S9; see *Methods*).

**Figure 3:**
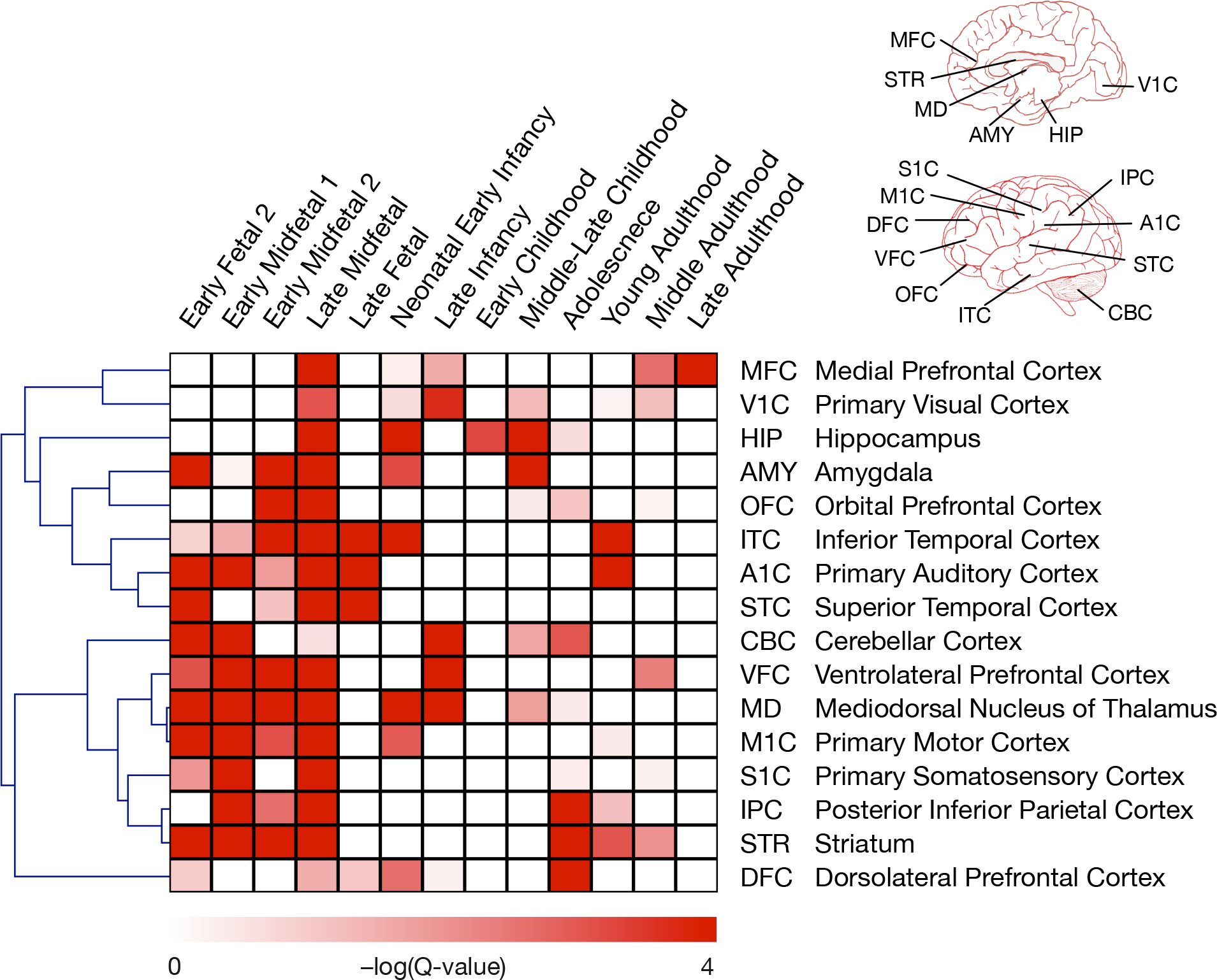
ASD-associated genetic changes in the spatiotemporal development of the brain. The heatmap shows the enrichment of spatiotemporal gene expression signatures towards the top of the genome-wide ranking of ASD genes. The 16 brain regions and 13 developmental stages considered labelthe rows and columns, respectively. The regions are further marked on illustrations of the human brain at the top right. Each cell in the heatmap (row, column) corresponds to a spatiotemporal signature-a set of genes highly expressed specifically in that region (row) at that developmental stage (column). The intensity of the color in each cell (scale below) representsthe log-transformed significance of ASD-association of that signature. A rank-based scoring followed by a permutation test was used to calculate *P* values, which were then converted to *Q* values to account for multiple hypothesis testing (see *Methods*). The heatmap shows a striking prenatal signal suggesting major effect of ASD-associated mutations on fetal brain development.

Our analysis identifies a clear developmental pattern-a prenatal signal from the early, mid, and late fetal stages-indicating that autism-associated genetic changes affect the development of the fetal prefrontal, temporal, and cerebellar cortex. This finding is largely consistent with previous studies^17, 20, 40, 45^ (reviewed in^46^), as well as recent findings on the possible role of callosal projection neurons^19^ and on the neuropathology of autism brain^47^. Spatially, on the other hand, we observe that the activity of ASD candidate genes is widely distributed across regions during early brain development, displaying a broad distribution that underscores the heterogeneity of the disorder. This pattern is consistent both with the lack of a single, highly reproducible neuroanatomical correlate of autism^48^, and with evidence that autism may be a disorder of general neural processing that manifests in many regions, including visual, auditory, motor, and somatosensory cortices^49^.

Performing this analysis using our genome-wide ASD candidate genes not only aids in revealing high-level spatiotemporal patterns relevant to ASD,but also enables us to now localize hundreds of novel candidate genes to specific brain regions that have been previously linked to the etiology of autism, such as the cerebellum^50^ (reviewed in^51^) and striatum^52 53^. A substantial body of literature has implicated the cerebellum in the development of the disorder, both in mouse models^50^ and in humans (reviewed in^51^), and our results highlight both fetal and early postnatal time points when disrupted gene expression in the cerebellum may contribute to autism risk. In addition, the striatum is identified here as a key locus of autism pathology throughout fetal development, a finding of particular relevance to repetitive behaviors as explored in mouse models that have both construct and facevalidity^52, 53^. In addition to known ASD-linked regions, we observe a new molecular signal in the mediodorsal nucleus of the thalamus in early and mid-fetal brain development that is consistent with reduced functional connectivity of this region in autism as suggested by fMRI^54^. Together, our analysis provides a way for choosing specific regions at relevant developmental stages for experimental verification of novel ASD candidate genes at the highest spatiotemporal resolution allowed by the expression data.

### Brain-specific functional modules disrupted in ASD

In concert with studying the developmental aspects of autism, a detailed understanding of the underlying effects of autism-associated genetic changes on cellular functions and pathways will result in a more complete picture of the molecular basis of ASD. Typical pathway analysis of known ASD-associated genes falls short of this goal, with the ability to only highlight functionally annotated genes, while providing little information about the hundreds of uncharacterized genes, let alone how all these genes and associated pathways are functionally interconnected in the brain. The humanbrain-specific functional gene interaction network that underlies our autism gene predictions lends itself naturally towards addressing this challenge. First, we define an autism-associated network by restricting the genome-wide brain network to the top 10% (∼2,500) of the predicted ASD-associated genes. Then, we use a strategy based on shared k-nearest-neighbors (SKNN) and the Louvain community-finding algorithm^55^ to cluster and partition the network into distinct and significantly cohesive functional modules predicted by our approach to be affected by ASD-associated genetic changes (Figure 4, Figure S10; see *Methods*). Based on the intuition that genes that functionally cluster together tend to have common local neighbors in the network^56^, the SKNN method calculates the number of shared nearest neighbors between ASD candidate genes in the brain network. The Louvain algorithm then finds communities of ASD candidates with shared local neighborhoods. Thus, the SKNN-based strategy has the advantages of alleviating the effect of high-degree genes and accentuating local network structure by connecting genes that are likely to be functionally clustered together.

**Figure 4:**
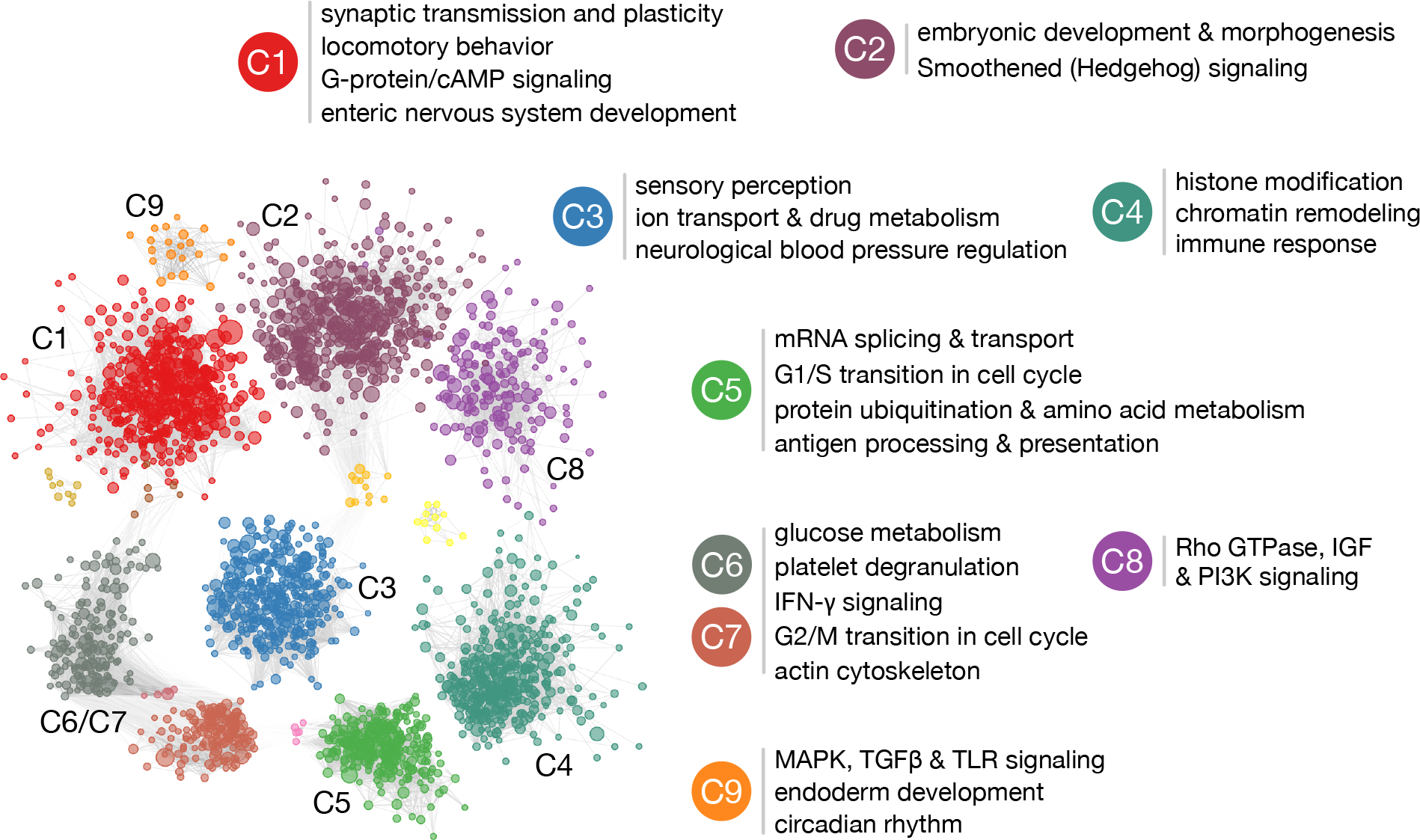
Autism-associated brain-specific functional modules. The network of brain-specific functional interactions among the top 2,500 ASD-associated genes were clustered using a shared-nearest-neighbor-based community-finding algorithm (see *Methods*) to elucidate several modules of genes (left). Nine of the clusters that contained 10or more genes, labeled C1 through C9, were tested for functional enrichment using genes annotated to Gene Ontology biological process terms. Representative processes/pathways enriched within each cluster are presented here alongside the cluster label. Since C6 and C7 shared a number of strong links across the clusters, they were merged before calculating functional enrichment. The enriched functions provide a landscape of cellular functions potentially dysregulated by ASD-associated mutations.

The primary contribution of this analysis is a systematic way of associating individual genes-several among them being novel and uncharacterized-to specific processes and phenotypes that might underlie autism etiology (Figure 4, Table S6). This includes cellular functions (e.g., synaptic transmission and neuronal function^8, 12, 31, 57^ (cluster 1, C1)), signaling pathways (e.g., canonical Wnt^4, 21^ (C2), gamma interferon^58^ (C6 and C7),), and fundamental cell biological processes (e.g. histone modification and chromatin remodeling^7, 8^ (C4) and cell cycle regulation (C5)) that have been repeatedly implicated in autism. The clusters delineated here also connect genes with key autism-associated phenotypes. For example, genes involved in enteric nervous system development (C1) might underlie the gastrointestinal symptoms common and persistent in children with autism^59^. Similarly, hypersensitivity to a range of stimuli is a well-established aspect of the autism phenotype^60^, and the genes identified in C3 include those involved in a whole assortment of perceptions (e.g., chemical stimuli,sound, light, smell, pain, temperature). Also noteworthy given a strong association of sleep dysregulation with autism^61^, is circadian rhythm captured in C9.

These findings suggest functional roles for genes associated with ASD pathogenesis and provide a framework to better understand the genes that underlie the emergence of important and often poorly understood phenotypes in animal models. Approximately 20% of the genes within each cluster are completely uncharacterized (lack any annotation to specific processes/pathways in the Gene Ontology), and this analysis suggests specific functional roles for these genes. For example, DIP2C (#180) and RSBN1 (#85) are top-ranked uncharacterized genes in C8 and C4, respectively,that are also specifically expressed during early fetal development. The grouping of DIP2C in C8, its fetal gene expression, and the identification of a *de novo* LGD mutation in DIP2C in an autistic proband in the Simons Simplex Collection^6^ together link DIP2C to autism through the Rho and IGFR pathways. Likewise, RSBN1 is tightly linked to chromatin remodeling and histone modification genes in C4. Its mouse homolog exhibits a punctate nuclear location^62^, consistent with a nuclear function for the protein. These and many other uncharacterized genes (Table S6) are candidates for experimental follow-up in specific functional contexts to characterize the cellular dysregulation in autism.

### Identifying ASD-associated genes within recurrent autism-associated CNVs

Copy number variations (CNVs) are important risk factors for autism. However, identifying which among the genes in these chromosomal intervals are pathogenic remains challenging. Our genome-wide prediction enables us to systematically prioritize genes within CNVs and reveal candidates for experimental verification. We applied this approach to eight of the most common autism-associated CNVs: 16p11.2, 15q11-13, 15q13.3, 1q21.1, 22q11, 7q11.23, 17q12, 3q29 (Figure 5). Further delving into the candidate genes, we garnered existing functional and genetic evidence and provide hypotheses about how perturbations of genes from different CNVs might converge at the molecular level by affecting specific biological pathways, leading to ASD.

**Figure 5:**
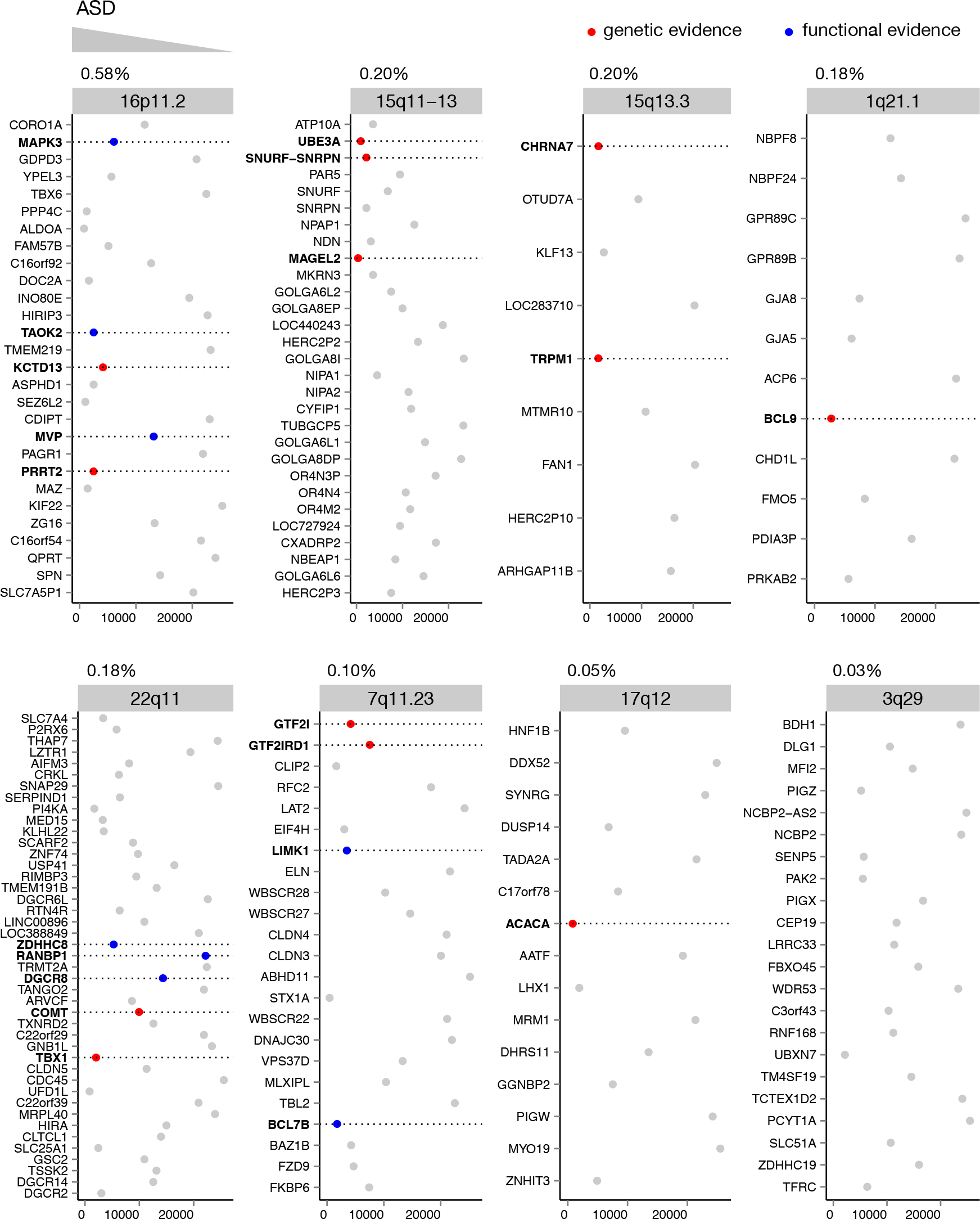
Prioritization of genes within eight recurrent ASD-associated CNVs. Each plot corresponds to one of the eight CNVs orderedbased on their observed frequency in autistic cases^15^ (given above each plot). The plots show the genes in each CNV interval in their genomic order (y-axis) and each gene’s ASD-association rank in our genome-wide predictions (x-axis) from top (low) ranks on the left to bottom (high) ranks on the right. The points corresponding to the genes are also colored red and blue based on whether there exists previously known (direct) genetic or (indirect) functional links between the genes and autism, independently curated by an ASD expert. The detailed ranking and evidence for CNV genes are in Table S7. The dashed lines are visual aids to read the gene names (in bold) of the colored point. Across the CNVs, genes with independent genetic evidence (red) and those with functional evidence (blue) tend to be ranked near the top of our genome-wide predictions (towards the left of each plot), compared to allthe other genes that lack any evidence (grey).

First, examining the genes that we prioritize within each of the eight autism-associated CNVs (genes within a CNV that are ranked in the first decile of our genome-wide ASD-gene ranking), we find that our network-based approach pinpoints many genes with previous genetic or functional evidence linking them to autism (Figure 5, Table S7). Across the CNVs, genes with prior genetic evidence (strongest, direct) are ranked significantly higher than those with functional evidence (indirect) (one-sided Wilcoxon rank-sum test *P* = 0.012), which are significantly ranked higher than CNV genes with no evidence of ASD association (*P* = 0.04; ‘genetic’ ranked higher than ‘no evidence’ *P* = 1e-6; Figure S11). Our method also implicates novel ASD candidates within these CNVs, which, based on the above trend, are strong candidates for further experimental follow-up. Experiments may particularly be a priority in cases where known functions of a highly ranked gene dovetail with biological processes that have already been implicated in the downstream effects of these CNVs. For example, a recent report suggests that changes in copy number at the 16p11.2 locus-the most widely studied and one of the most common CNVs linked to autism^63^-may affect RhoA signaling and dynamics of the actin cytoskeleton^64^. One of our top-ranked genes in 16p11.2 is PPP4C (ranked third within the interval), which is known to regulate Rho GTPases and levels of filamentous actin^65^, suggesting a mechanism whereby haploinsufficiency for PPP4C might contribute to the phenotypes associated with 16p11.2 deletion. Another potential link between 16p11.2 and ASD is the highly ranked transcription factor MAZ (ranked fourth within the interval; see *Discussion*). Strikingly, 10 out of the 23 top-predicted (first decile) genes across the eight CNVs, including MAZ, are significantly highly expressed in the mediodorsal nucleus of the thalamus (MD) during early-midfetal development, a spatiotemporal window that we associate with autism in this study (Fisher’s exact test *P* < 1.2e-9).

With multiple CNVs and a number of candidate genes within each, a further challenge is to identify their convergence on similar cellular functions that might be dysregulated to result in autism. Here, we attempt to shed light on this mechanism by leveraging the top-ranked genes within the CNVs and the brain-specific network to identify pathways and cellular processes that may be disrupted by ASD-associated CNVs. Intuitively, we hypothesize that genes involved in these disrupted pathways (referred to as ‘intermediate genes’) would be critical bridges between ASD candidate genes within CNVs and the major autism phenotype genes within the brain functional network. Practically, we identified such intermediate genes as those along the shortest paths in the brain network specifically linking the predicted ASD-associated genes within each CNV to the 19 high-confidence (E1) autism genes^25, 26^ that represent the core molecular etiology of autism (illustrated in Figure S12; see *Methods*). The intermediate genes we identified thus likely represent cellular functions that CNVs impinge on to contribute to the autism phenotype. We then outlined the biological processes enriched in these intermediate genes that were also co-mentioned with autism in the literature (see *Methods*). Through this analysis, we were able to propose, in a data-driven way, genes and processes that underlie the functional connection of ASD-associated CNVs with the ASD molecular phenotype (Figure 6, Table S8).

**Figure 6:**
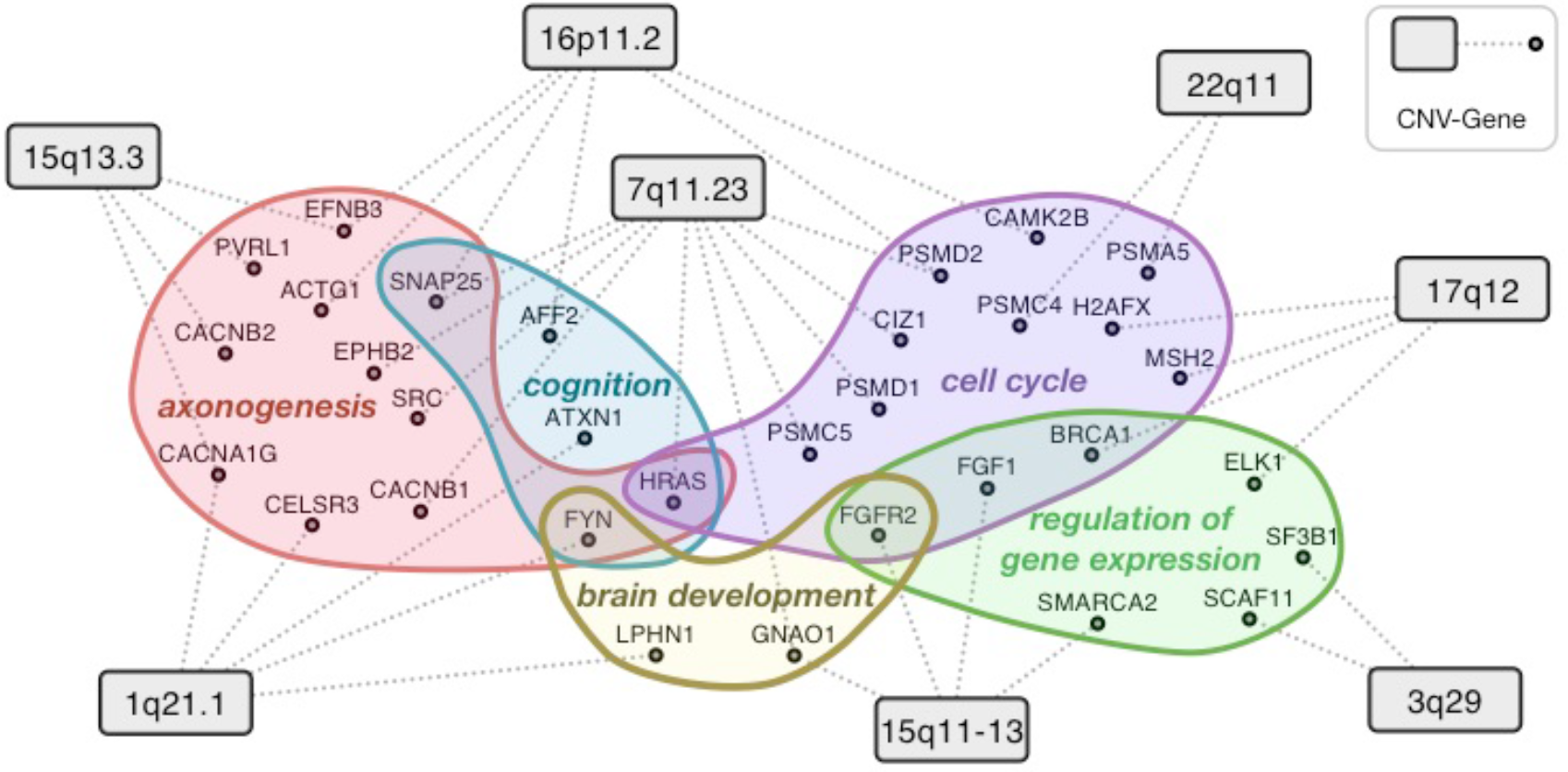
Convergence of cellular functions disrupted by multiple CNVs identified through key intermediate genes in the brain network. The network diagram above illustrates the intermediate genes linking the eight CNVs (black rectangles) to the molecular phenotype of ASD. The dotted grey lines represent high confidence functional links in the brain that mediate the linkage of top CNV genes’ to ASD genes; these linkages go through key intermediate genes (black circles, see *Methods*). Enrichment analysis groups the intermediate genes into a small number of autism related processes (illustrated as colored clouds). For visual clarity, only representative examples of processes associated with at least two CNVs are included. The functions of these intermediate genes illustrate the hypothesis that multiple CNVs might disrupt a core group of ASD-related biological processes and pathways.

Multiple previous studies support our findings. For example, SNAP25, one of the intermediate genes mediating 16p11.2’s linkage with cognition/axonogenesis, is reported to bind PRRT2 in 16p11.2 to regulate neuronal processes^66^. Besides these well-established CNV-gene-process linkages, many intermediate genes identified are themselves associated with autism. SMARCA2, which mediates 15q 11-13’s association with regulation of transcription, is part ofthe BAF complex and is implicated in autism^67^. AFF2, linked with 16p11.2, is identified in multiple autism studies with rare mutations^68, 69^. EPHB2, linking 7q11.23 with axonogenesis, harbors two *de novo* variants in ASD probands in separate sequencing studies. LPHN1-linked to 1q21.1-and SF3B1-linked to 3q29-were identified recently to be among 239 genes most likely involved in ASD based on *de novo* mutation and absence of mutation in healthy controls^34^. CACNB2, linked to 15q13.3, was recently added to category 3in the SFARI Gene database^25^. We also noticed that all three genes linking 17q12 to cell cycle (BRCA1, MSH2, H2AFX) are either directly or indirectly involved in DNA damage repair, which was highlighted in a postmortem gene expression study as an autism-associated process^70^. This observation is in agreement with the hypothesis that dysregulation of mismatch repair might contribute to the generation of somatic mutations in the brain that contribute to autism risk. The full set of intermediate genes and enriched processes (Table S8) provide a rich resource for experimental follow-up studies.

### A web-interface to explore autism-gene predictions

We have developed a dynamic web interface-asd.princeton.edu-that provides an interactive portal to all autism-gene predictions from this study along with the results from the subsequent analyses, including spatiotemporal brain signatures, functional modules, and prioritized CNVs. Using this interface, autism researchers can explore the results from this study not just as ranked genes, but also their inter-connections in the context of the human brain-specific network. The interface allows gene-based search, dynamic network visualization, and ability to export results (Figure S13).

## Discussion

The genetic heterogeneity underlying ASD is daunting, with multiple modes of inheritance, a sizeable contribution from *de novo* events, and hundreds of both rare and commonvariants likely contributing tothe disease^71^. Further compounded by diverse clinical phenotypes and limited sample sizes, it is unlikely that all risk genes would be identified by sequencing studies and statistical association alone. Sequencing screens mainly identify mutations with large effect size, and statistical quantitative genetic studies rely on relatively high mutation frequency, both of which are missing in gene variants with modest frequency and effect size. Moreover, all these methods fail to model epistatic effects that could underlie the ASD phenotype.

In this study, we present a powerful, genome-wide computational approach to facilitate ASD gene discovery. By combining evidence from thousands of gene expression, protein-protein interaction, transcription regulation and genetic perturbation data, we recently built a gene network that serves as a robust scaffold of brain-specific gene interactions and cellular functions^24^. Here, we take advantage of this brain-specific functional interaction network to provide a ranking of all genes in the human genome by their potential pathogenic involvement in ASD (Figure 1). The validity of this comprehensive complement of genes is attested by an independent exome sequencing study published after our initial analyses were completed (and, therefore, not included in our training sets), with the strong autism candidates from the study (recurrent *de novo* LGD mutations) ranking near the top of our prioritized list (Figure 2). Therefore, our ranking can, for example, be used to prioritize novel autism-risk genes for experimental functional characterization or choose gene candidates for targeted sequencing in new studies, thus increasing specificity and reducing cost. Our predictions can further help guide the analysis of whole exome/genome sequencing results, by prioritizing which genes harboring missense mutations or nearby noncoding variants should be followed-up by downstream experimental investigation.

Traditional disease-gene prediction methods^72^ use only the most-confident genes or all genes irrespective of the evidence linking them to disease. Our evidence-weighted machine learning method, which we used to make these genome-wide ASD candidate gene predictions, brings new prospects for incorporating a large number of genes with various levels of confidence of autism-association identified from multiple experimental/sequencing efforts. Even within a single study, different types of variants-LGD, missense, synonymous-potentially confer variable amounts of risk. Our weighted approach embraces this diffuse knowledge of disease genetics, and we demonstrate that our method is able to identify ASD-related signals among the noisy and weak gene associations present in the literature and make more accurate predictions (Figure 2a). Because the method is highly generalizable, it can be readily applied to the study of many other complex diseases.

In addition to prioritizing hundreds of novel ASD candidate genes, we have used the underlying network to link individual ASD genes to perturbed cellular functions and higher-level phenotypes (such as brain development,sensory perception, and circadian rhythm), thus aiming to bridge the genotype-phenotype gap in autism. In fact, our genome-wide prediction ranks LGD genes in SSC probands with lower non-verbal IQ (<100) significantly higher than LGD genes in probands with higher non-verbal IQ (> = 100; one-sided Wilcoxon rank-sum test *P* = 0.013). Similarly, LGD genes highly ranked by our method are also associated with severe restricted repetitive behavior^73^ (permutation test *P* = 0.02). Usinga combination of our whole-genome ranking of ASD candidate genes and spatiotemporal gene expression data of the human brain^44^, we have gained insights into specific stages in human brain development that might be critically affected by mutations in autism-associated genes. We provide the functional and developmental contexts of our top-decile predictions in Table S9. Furthermore, this analysis showcases how biomedical researchers can use our genome-wide ASD candidate gene predictions as a framework for analyzing their data from high-or low-throughput assays, allowing high-resolution study of autism genetics in the functional and physiological contexts of their interest. For example, a researcher who has generated gene expression or proteomics data from anew high-resolution spatiotemporal window in the brain (either human or ina model organism) can use our approach to assess the molecular relevance of that window to ASD. This approach-identifying a characteristic gene signature from the samples and estimating its enrichment in our genome-wide ASD-gene ranking (similar to results presented in Figure 3)-can provide results of increasing specificity with increasing spatiotemporal resolution in the available data. (S)he could then focus on the highly-ranked genes in her/his expression signature in combination with the specific functional contexts identified for these genes (‘ASD-associated functional modules’ as presented in Figure 4) to generate hypotheses and design experiments to further characterize autism.

Finally, we have used the genome-wide predictions to prioritize potentially “causal” genes within large regions of autism-associated CNVs and illustrate how genes from different CNVs converge onto a set of common genes and pathways that could mediate the cellular perturbations associated with autism. This analysis provides experimentally-testable hypotheses for further research into the molecular mechanisms underlying the connection to ASD for the most common ASD-associated CNVs. For example, one of the top ranked ASD candidate genes in 16p11.2 is the transcription factor MAZ (ranked fourth in the 16p11.2 interval), which has not been previously characterized in the context of autism. On the genome-scale, computationally predicted targets of MAZ (genes with MAZ’s DNA-binding motif^74^ in their upstream sequences) are statistically over-represented in our ASD-associated gene predictions (*P* = 2.5e-93), and are involved in the regulation of neuron differentiation and axonogenesis. MAZ and its predicted targets also show specific spatiotemporal expression in the early mid-fetalstage of development in the primary motor cortex (M1C; Figure 3), a stage demonstrated to be relevant to autism^40^. Together these observations lead to a specific hypothesis: the disruption of normal MAZ function (or one of its targets) in the primary motor cortex at the early mid-fetal developmental stage can lead to the autism phenotype by perturbing the transcription of neuronal differentiation and neurodevelopmental genes.

In addition to specific analyses and hypotheses that we describe in this study, we have made all the genome-wide predictions, ASD-associated brain developmental stages, functional modules, and prioritized CNVs from thisstudy, as well as their brain network contexts, available to the biomedical community via a dynamic web interface at asd.princeton.edu. Researchers can explore the top predicted candidate genes and processes as well as any genes they are specifically interested in (such as genes found in sequencing or GWAS studies) in the context of our network and generate new sets of related genes or processes. The top-ranked genes identified here, especially the many novel uncharacterized genes, have the potential to speed up gene discovery and spur further focused experimental studies that can lead to better objective genetic biomarkers for early diagnosis of ASD. We also expect that these genes and associated pathways, along with their brain-specific functional contexts, will help identify potential therapeutic targets that can make intervention more impactful. The predictions and analysis in asd.princeton.edu will be updated regularly as new genetic/sequencing studies of ASD are published to keep the results of this work relevant in the longer run.

## Methods

### Sources of known autism-associated genes

Several databases contained information about a number of autism-associated genes. Each resource, however, gathered data from sources with different levels of evidence, ranging from recurrent mutations in autistic patients to nebulous links gleaned from text-mining thousands of PubMed abstracts. In an effort to be comprehensive in putting together our initial picture of autism genetics, we collected genes linked to autism from all these sources (up to Dec. 2013) and designated them to four evidence levels, each associated with an evidence-weight that was commensurate with the quality of the evidence (Table S1). For example, genes from categories 1 and 2 in SFARI^25^ found by statistically significant rare variant associations or nominally significant (but replicated) common variant associations, as well as genes from OMIM^26^ (a total of 19 genes) were designated as evidence-level 1 (E1) and given a weight of 1.00. 31 candidate genes with two lines of literature evidence annotated to category 3 in SFARI were designated as evidence-level 2 (E2) and given a weight of 0.50. Databases HUGE^75^ and GAD^76^ together record 413 genes identified based on genetic association studies, which were designated to evidence-level 3 (E3). 131 genes inferred purely based on text-mining PubMed abstracts from the Gene2MeSH (gene2mesh.ncibi.org) and DGA^27^ databases, as well as those assigned with minimal-evidence (to category 4) in SFARI were designated as evidence-level 4 (E4). Genes in levels 3 and 4 were given a weight of 0.25. Each gene was uniquely assigned to the highest evidence-level. In total, we curated 594 autism-associated genes with evidence-weights ranging from 0.25 to 1.

### Human brain-specific functional interaction network

The human brain-specific network is a scaffold of predicted functional interactions between all pairs of genes in the genome. Each edge (link) inthe network between a pair of genes is associated with a probability that represents how likely the two genes are to participate in the same biological process specifically in the brain. This network is the product of systematic integration of thousands of genomic datasets in the context of known functional gene interactions in the brain^24^ (Figure S13). We have shown that multiple genes associated with a particular disease often tend to have similar patterns of connectivity in a relevant tissue-specific functional network^24^. We use this notion (further described below) to discover autism-associated genes in the context of the brain-specific gene network.

### Learning and cross-validation of the network-based classifier

We use a machine learning approach to predict autism-association genes.Our approach has two steps: 1) Building a statistical model that captures the connectivity patterns of known autism genes in the brain-specific network, and 2) Using this model to subsequently predict whether each of the other “unknown/unlabeled” genes in the network looks like an autism gene based on its connectivity in the network. We trained an evidence-weighted linear support vector machine (SVM) classifier using the gold standard of known autism-associated genes (along with their weights) as positives and 1189 genes associated with non-mental-health diseases (from OMIM, with weights equal to 1) as negatives. The brain-specific interaction probabilities of each gene to all the genes in the brain network were used as features. Given the network features (*x_i_*) and a positive/negative label (*_y_*) for all the *m* training genes, the linear SVM solves the following optimization problem^77^:

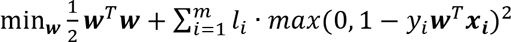
where *l_i_* is a penalty parameter specific to each gene that influences how costly it is to misclassify that gene. By setting *l_i_* equal to the evidence-weight of the labeled gene (where negatives have *l_i_* of 1), we ensure that we train a model that rarely misclassifies high-confidence genes while giving certain latitude in correctly classifying low-confidence genes.

To evaluate this approach, we employed a stringent five-fold cross-validation scheme: in each fold, we trained a model on 4/5th of the labeled (positive and negative) genes and evaluated the model on only high-confidence (E1) positives (and all negatives) in the remaining 1/5th of the labeled genes. The model with weights equal to 1.00, 0.50, 0.25, and 0.25 for E1, E2, E3, and E4 genes had an area under the receiver-operator-curve (AUC) of 0.80. Taking model variance into account by repeating 5-fold cross-validation 50 times, we demonstrate that this model performs significantly better than one trained only using high-confidence genes (weights 1, 0, 0, 0; AUC = 0.73; *P* < 2.2e-16, one-sided Wilcox rank based test; weights 1, 1, 0, 0; AUC = 0.76; *P* < 2.4e-15), and one trained with real E1-2 supplemented with random genes simultaneously matched with E3-4 for gene length, brain expression, and neuronal gene ontology (AUC = 0.75, *P* < 2.2e-16). Therefore, we used the evidence-weighted model further to make predictions.

### Genome-wide prediction using evidence-weighted network classifier

We coupled whole-genome prediction with 5-fold cross-validation: a prediction for each labeled gene (positive/negative) was recorded only from the fold that did not include the gene for training, thereby relying only on its network-based similarity to other genes instead of its own prior evidence; a prediction for an unlabeled gene was recorded as the average of predictions from the five folds. Each prediction corresponded to the distance of the gene from the hyperplane that separates positive genes from negative genes, and these distances were used to rank all the genes in the genome.For interpretability, the distances were also converted to probabilities using isotonic regression (see below). This produced a genome-wide ranking of 25,825 genes based on their predicted level of association with autism.

### Evaluation of autism gene ranking on independent *de novo* mutations

#### De novo mutations from exome-sequencing study

The Simons Simplex Collection^78^ (SSC) contains more than 2,500 families, each of which has a single child with ASD (referred to as proband). Most of these families have at least one unaffected sibling. A recent study used whole exome sequencing of the SSC to identify *de novo* LGD mutations in autistic children and their unaffected siblings^7^. 350 and 174 genes were targets of LGD mutations in autistic probands and unaffected siblings, respectively. Among the 350 LGD genes in probands, 27 were observed in more than one proband (recurrent LGD), indicating that these are very likely to be true autism-associated genes. Weused these sets of genesproband recurrent LGD (27), proband LGD (350) and sibling LGD (174)to benchmark our genome-wide autism gene ranking. When testing each LGD set, we used 462 genes containing synonymous mutations in unaffected siblings (sibling SYN) as a control. Further, to account for potential biases in exome sequencing coverage, when defining a genomic background, we removed the 8,054 genes in which no rare variants from reference were reliably detected anywhere within genic boundaries.

Since preliminary *de novo* mutation data on 682 SSC families were published in three separate reports in 2012^3, 4, 6^, and it was likely that some of these data were represented in our training set of autism-associated genes, we created an “unpublished” evaluation data set from SSC which was released after our prediction. This data set was created from the new SSC families in 2014 study^7^ (i.e., 1,835 families after removing the previously published 682 from the total of 2,517 families), which performed whole exome sequencing to identify de novo LGD mutations in autistic children and their unaffected siblings. We used this “unpublished” set of mutations to retestthe significance of our findings derived based on the entire SSC cohort (Figure S4).

We also obtained the 107 likely ASD-associated genes from a study independent of the SSC^8^ and used this set to evaluate our predictions.

#### Decile enrichment test

After removing SSC genes without rare variants from the autism gene prediction rankings, the remaining genome-wide gene ranking was first divided into deciles. For a given set of LGD genes, we used the binomial test to assess whether a larger fraction of the LGD genes occurred in the first decile when compared to the expected fraction based on the occurrence of sibling SYN genes. Reassuringly, sibling SYN genes are roughly apportioned 10% to each decile, indicating that it is a good control set.

#### Rank-based enrichment test

To demonstrate robustness of our evaluation to specific rank cutoffs (as our decile enrichment test assesses genes above a 10% rank cutoff), we also formulated a rank-based test. This test takes the entire genome-wide ranking and assesses if the set of LGD genes have a skewed distribution towards the top of the ranking relative to the control set. First, based on the genome-wide autism-association ranking, we calculated an exponential score *s_i_* for each gene *i* based on its autism-associated rank *rank(i)* as follows:

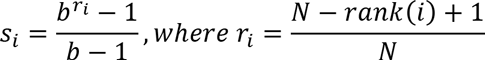

*N* is the total number of genes equal to 25,825, and *b* was set to 100. The score s_i_ ranges between 1 for the top-ranked gene dropping exponentially to 0 for the lowest ranked gene. The rank-based test then proceeds in three steps: (1) calculate the observed difference (*d_obs_*) between the mean rank-based score of the test LGD gene set (e.g. proband LGD) and the control gene set (sibling SYN); (2) shuffle the gene labels in the test and control gene sets 100,000 times and, each time, record the difference between *mean(test)* and mean(control) (*d_permut_*); (3) calculate a *P* value for the observed difference (*d_obs_*) equal to the fraction of permuted differences *d_permut_* that were equal to or greater than *d_obs_*; also calculate an effect size for *d_obs_* as a z-score:

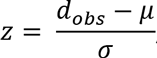
whereμ and σ are the mean and standard deviation of the distribution of *d_permut_* values.

#### Evaluation controlling for gene length

Genes linked to ASD are known to be longer in size than genomic background^29^. To test if our genome-wide ranking is driven by prioritizing long genes, we repeated our LGD evaluation while controlling for gene length. We sorted all genes by their length, divided the genes into bins of 500 genes (of very similar lengths), randomized ASD ranks of genes within each bin, and calculated the top-decile enrichment of LGD-target genes. A *P* value was calculating by repeating this procedure 10,000 times and calculating the fraction of times the permuted top-decile enrichment *P* value was less than or equal to the observed *P* value.

#### Evaluation controlling for bias in brain-/neural-functional annotations

To demonstrate that brain/neural genes are not biasing our results, we evaluated our predictions on whether they can prioritize proband LGD genes within a non-neural-annotated gene set. We created this set by including proband LGD genes that are neither annotated to neural/brain-related functions in Gene Ontology nor part of our training gold standard. Fisher’s exact test was then used to calculate significance.

#### Evolutionary constraint of ASD-associated genes

Evolutionary constraint estimates of genes were obtained from two different studies^79, 80^, one providing a quantitative (RVIS) score for each gene, and the other providing a single set of 1,003 constrained genes. The Wilcoxon rank-sum test was used to assess the difference between the RVIS scores for genes in our top-decile compared to the rest of the ranking, and the Fisher’s exact test was used to calculate the enrichment of the constrained gene set in our top decile. We also evaluated both sets using a rank-based permutation test that does not involve any rank cutoffs. For the RVIS score, a test statistic equal to the Spearman correlation coefficient between genes ranked by our method and by RVIS scores was calculated; a *P* value for this statistic was estimated by permuting the ASD candidate gene ranking a million times and calculating the fraction of times the permuted correlation was less than or equal to the observed one. For the constrained genes, a test statistic equal to the mean rank-based scores (*si*) of genes in the set was calculated; a *P* value for this statistic was then estimated by permuting ASD gene ranks a million times and calculating the fraction of times the permuted statistic was greater than or equal to the observed one.

### Developmental expression of autism-associated genes in the brain

#### Defining spatiotemporal expression signatures

The spatiotemporal developmental gene-expression data for the human brain was obtained from Brainspan^44^. Raw data from this study were downloaded from NCBI GEO^81^ accession GSE25219. CEL files were background corrected, normalized, and summarized using RMA^82^ based on a custom CDF^83^, and expression for each gene was averaged across replicates. Further analysis was restricted to 13 stages from early fetal (10-13 weeks post-conception) to late adulthood (≥60 years) that contained expression data for all 16 brain-regions. We first established a gene-expression signature for each of the 208 spatiotemporal windows (combination of 16 regions and 13 stages) that was specific to that window with respect to other stages as well as other regions. For example, a signature for the striatum (region) at the late midfetal stage, was calculated in three steps:

1. For each gene *i* two modified z-scores were calculated by comparing *e_i_^rs^*, the gene’s expression in that region (striatum) and stage (late-midfetal), to the distribution of its expression values across i) all regions at late-midfetal stage (to get *z_i_^region^*., and ii) all stages of striatum (to get *z_i_^stage^*

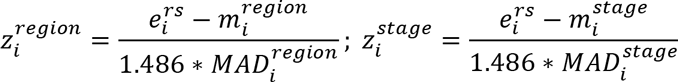
where *m_i_^region^*. and *m_i_^stage^*, respectively, are the median expression levels of gene *i* across all regions and all stages; *MAD_i_^region^* and *MAD_i_^stage^* are the corresponding median absolute deviations (MAD). Since median and MAD are stable measures of central tendency and variance of a distribution without being influenced by outliers, scaling using these measures aids in identifying expression values that are particularly high.
2. These two z-scores were then combined into a meta z-score *z_i_^rs^* that tends to be high when the expression of the gene deviates substantially from its nominal expression along both the spatial and temporal axes.

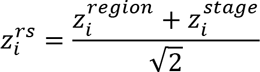
3. Finally, the set of genes with *z_i_^rs^*≥2 were used as the gene-expression signature (of late midfetal striatum, in this example).

We employed this procedure to compute a signature for all 208 combinations of regions and stages.

#### Rank-based enrichment test of spatiotemporal signatures

This test was very similar to the rank-based test detailed above. For each signature, a test statistic equal to the mean exponential rank-based autism-association scores (*s_i_*; described above) of genes in that signature was calculated. Mean scores based on random signatures of the same size sampled from the union of all genes in all signatures were then used to set up an empirical distribution. A *P* value for the signature was calculated ultimately as the fraction of permuted mean scores that were equal to or greater than the observed mean score for the real signature. We carried out this test for all 208 region-stage combinations and corrected for multiple tests using the Benjamini-Hochberg correction^84^ to get *Q* values.

### Clustering the autism-associated brain network

In order to identify autism-associated functional modules in the brain,we created a subset of the brain-specific functional network containing the top 2,500 autism-associated genes from our genome-wide ranking and all the edges between them. Then, we used an approach based on shared k-nearest-neighbors (SKNN) and the Louvain community-finding algorithm^55^ to cluster the network into distinct modules of tightly connected genes. Given a graph *G* with *V* nodes/genes and *E* edges, with each edge between genes *i* and *j* associated with a weight *p_ij_*, this technique proceeds the following way: (1) calculate a new weight for the edge between each pair of nodes *i* and *j* that is equal to the number of *k* nearest neighbors (based on the original weights p,y) shared by *i* and *j;* (2) choose the top 5% of the edges based on the new edge weights and apply a graph clustering algorithm. This approach has two key desirable characteristics: a) choosing top *k* instead of all edges deemphasizes high-degree ‘hub’ nodes and brings equal attention to highly specific edges between low-degree nodes; and b) emphasizing local network-structure by connecting nodes that share a number of local neighbors automatically links genes that are highly likely to be part of the same cluster. We used a *k* of 50 to obtain the shared-nearest-neighbor autism-associated brain network and used the Louvain algorithm to cluster this network into distinct modules. We confirmed that the node membership and cluster sizes are robust by testing a range of values for *k* from 10 to 100. To stabilize clustering across different runs of the Louvain algorithm, we ran the algorithm a 100 times and calculated cluster co-membership for all pairs of genes that was equal to the fraction of times (out of 100) the pair was assigned to the same cluster. These co-membership scores were used to layout the network in Figure 4 (using Cytoscape^85^), which represents all clusters that contained at least 10 genes, defined by a co-membership score≥0.9. We confirmed the statistical significance of this score using a permutation test, randomizing the k-nearest neighbors of each node in the network of top ASD-genes and redoing the clustering procedure (Figure S8).

One-sided Fisher’s exact test was used to find GO biological processes^86^ enriched in each cluster. Benjamini-Hochberg correction was used to correct for multiple tests and GO terms with *Q*≤0.1 were deemed significant. The entire table of enriched GO terms is provided in Table S6.

### Prioritizing genes in autism-associated CNVs

We selected the eight most statistically significant and frequent autism-associated CNVs^15^ and obtained the genes in the intervals from UCSC^87^. An expert independently annotated genes in each interval with genetic or functional evidence for association with autism from existing literature. Genetic evidence refers to direct genetic evidence implicating a gene in autism or a related disorder (e.g., schizophrenia, epilepsy). Functional evidence refers to the annotation of a gene to a function, process, or pathway involved in autism, but without direct genetic evidence.

#### Identification of CNV-specific intermediate genes

The goal here was to identify key genes and the related cellular functions that may be deregulated by each CNV by tracing a biomolecular path from the CNV to the molecular phenotype of autism. We first marked the most autism-associated genes in each CNV as those within the top 10% of our genome-wide autism-associated gene ranking. Then, we defined the 19 E1 (high-confidence) genes as the core genes representing the molecular phenotype of autism. Taking advantage of the underlying brain-specific network,for each CNV, we calculated the betweenness centrality (BC) of each network gene as the fraction of shortest paths from the top-ranked CNV genes to the core autism genes that also pass through that gene. The brain network was pre-filtered to contain only around top 1% of its edges, and path length was calculated as the reciprocal of the functional linkage score. Genes with high BC are molecular mediators between the CNV and autism. We identified genes with high BC that were also specific to the given CNV (with *T* total genes and *t* top-ranked genes) by keeping the core autism genes constant and repeating the BC calculation for random set of *t* genes from random genomic intervals with *T* genes. Finally, a permutation-based *P* value was computed for each network gene as the fraction of times that gene’s BC with random intervals was equal to or greater than that with the real CNV (n = 100,000). Upon Benjamini-Hochberg correction for testing the BC of thousands of genes, those with *Q*≤0.1 were identified as mediator genes specific to the given CNV.

#### Text mining for autism-associated processes

We queried all available PubMed abstracts from 2000 to May 2015 for (autism OR ASD OR autistic), and retrieved these autism-associated abstracts.GO biological processes terms were mapped to each abstract using simple text search, and a given term was declared as associated with autism if it was found in at least two abstracts.

#### Functional impact of multiple CNVs

The next goal was to use the CNV-specific mediator genes to determine cellular functions that are likely deregulated by multiple CNVs. For this, we used the Fisher’s exact test to find sets of genes annotated to autism-associated GO biological processes that overlapped significantly with the CNV-specific mediator gene set. Processes/pathways that were significantly associated (*Q*≤0.05) with two or more CNVs,and annotated to at least two mediator genes were analyzed further and were summarized to general terms based on gene overlap, term description, and their relationships in the ontology.

#### Identification of MAZ targets

Computationally predicted targets were generated by using FIMO^88^ to map the binding site of MAZ (obtained from HOCOMOCO^74^) to the 1,000bp upstream regions of all the genes in the genome. 2,446 genes with a match at *Q*≤0.01 were selected as potential targets of MAZ. Enrichment analysis for MAZ targets against the autism-gene predictions was applied using the one-sided Fisher’s exact test against the top 2,500 ranked gene predictions (top decile).

### ASD webserver

The web interface is implemented using the D3 library^89^ for visualization, which enables use on any modern web browser without plugin installation.

#### Probability estimates for predicted autism-related genes

We estimated priors for several biological pathways and gene sets known to be enriched for autism-associated genes by calculating the fraction of genes with *de novo* LGD mutations occurring in quad family probands over all *de novo* LGDs. These pathways and sets include CHD8 targets, chromatin remodeling genes, FMRP targets, MAPK pathway genes, post-synaptic density genes, TOP1 targets and Wnt/beta-catenin pathway genes.

To improve interpretability for our autism predictions, we calibrated the SVM scores using isotonic regression. Isotonic regression, or monotonic regression, minimizes the same condition as a least-squares regression while imposing a monotonicity constraint (i.e., if gene A has a higher rank in original predictions relative to gene B, then isotonic regression enforces that the fitted probability estimate for gene A will also be higher than that of gene B), and it has been previously shown to have more power when sufficient data is available^90^. The response variable (probability of a gene being autism-related) was estimated based on their enrichment among the SSC proband LGDs. To prevent overfitting in our probability estimation, we used 10-fold cross validation to find the “knots” for the regression results of each held-out fold. We fit an isotonic regression curve through all knots,with hermite spline interpolation between knots, and provide the resulting estimated probabilities on the web interface.

#### Permutation-based P values and Q values for our genome-wide predictions

In order to improve the interpretability of our genome-wide ranking, we calculated a permutation-based *P* value and a corresponding *Q* value for each gene (Table S3), and provide these in our ASD web-server. We permuted labels of gold standards and retrained the model 1000 times. *P* values were calculated as the probability of the gene being predicted to be as extreme as it is (considering both ‘highly ASD-associated’ and ‘highly non-ASD associated’, *n* = 1, 000), and were corrected to *Q* values by the Benjamini-Hochberg method^84^.

#### Webserver updates

We will keep the ASD gene predictions on our web server updated for the community. As the set of annotated ASD candidate genes increases, we will retrain our model with the most up-to-date ASD candidate gene collection. We have already retrained our model with the entire collection of known ASD genes up to Oct. 2015 (with the same gene evidence weighting criteria), and those predictions are available at asd.princeton.edu/v2. We see that the new genome-wide gene ranking is largely consistent with the predictions used in this study, with a correlation of 0.93 in overall gene ranking and 83.3% of genes in the top decile matching the original predictions.

## Acknowledgements

We are grateful to all of the families at the participating Simons Simplex Collection (SSC) sites, as well as the SSC principal investigators.

We thank all members of the Troyanskaya Lab for valuable discussions. We thank John Spiro and other members of the Simons Foundation for constant feedback on the work and manuscript.

This work was primarily supported by US National Institutes of Health (NIH) grants R01 GM071966 and R01 HG005998 to O.G.T. V.Y. was supported in part by US NIH grant T32 HG003284. This work was supported in part by US NIH grant P50 GM071508. O.G.T. is a senior fellow of the Genetic Networks program of the Canadian Institute for Advanced Research (CIFAR).

